# KH domain containing RNA-binding proteins coordinate with microRNAs to regulate *Caenorhabditis elegans* development

**DOI:** 10.1101/2020.08.03.235127

**Authors:** D Haskell, A Zinovyeva

**Affiliations:** Division of Biology. Kansas State University. Manhattan, KS, 66506

**Keywords:** microRNA, RNA binding protein, KH domain, hnRNPK

## Abstract

microRNAs (miRNAs) and RNA binding proteins (RBPs) regulate gene expression at the post-transcriptional level, but the extent to which these key regulators of gene expression coordinate and the precise mechanisms of their coordination are not well understood. RNA binding proteins often have recognizable RNA binding domains that correlate with specific protein function. Recently, several RBPs containing K Homology (KH) RNA binding domains were shown to work with miRNAs to regulate gene expression, raising the possibility that KH domains may be important for coordinating with miRNA pathways in gene expression regulation. To ascertain whether additional KH domain proteins functionally interact with miRNAs during *Caenorhabditis elegans* development, we knocked down twenty-four genes encoding KH-domain proteins in several miRNA sensitized genetic backgrounds. Here, we report that a majority of the KH domain-containing genes genetically interact with multiple miRNAs and Argonaute *alg-1*. Interestingly, two KH domain genes, predicted splicing factors *sfa-1* and *asd-2*, genetically interacted with all of the miRNA mutants tested, while other KH domain genes exhibited functional interactions only with specific miRNAs. Our domain architecture and phylogenetic relationship analyses of the *C. elegans* KH domain-containing proteins revealed potential groups that may share both structure and function. Collectively, we show that many *C. elegans* KH domain RBPs functionally interact with miRNAs, suggesting direct or indirect coordination between these two classes of post-transcriptional gene expression regulators.

## INTRODUCTION

Most developmental and cellular processes rely on precise choreography of gene regulatory networks that incorporate a wide range of cellular and environmental inputs. Evolution of multiple regulatory pathways provided cells with multifaceted and combinatorial methods of regulating gene expression allowing for robustness, flexibility and rapid remodeling of expression patterns. One essential layer of gene regulation occurs at the post-transcriptional level and is effected by two classes of molecules: small non-coding RNAs called microRNAs (miRNAs) and RNA binding proteins (RBPs). The human genome is predicted to encode at least 2000 miRNAs (Alles *et al.* 2019) and approximately 1500 RBPs (Gerstberger *et al.* 2014). In comparison, *C. elegans* genome is predicted to encode more than 180 miRNAs (Ambros and Ruvkun 2018), and at least 850 RBPs (Tamburino *et al.* 2013) making it a more tractable model to study the functional interactions between miRNAs and RBPs.

Most mature miRNAs are generated via a canonical multi-step biogenesis pathway that starts with transcription of primary miRNA (pri-miRNA) transcripts. Pri-miRNAs are then processed by consecutive enzymatic activities of Drosha and Dicer endonucleases to generate a double stranded RNA duplex, which is ultimately loaded into an Argonaute protein. A single miRNA strand is retained by Argonaute and the mature miRNA silencing complex (miRISC) is formed when the miRNA-loaded Argonaute associates with a GW182 effector on the target messenger RNA (mRNA) (reviewed in Gebert and MacRae 2019). The miRISC identifies target mRNAs through partial sequence complementarity, ultimately resulting in translation repression and/or mRNA degradation (reviewed in O’Brien *et al.* 2018; Gebert and MacRae 2019).

RNA-binding proteins regulate diverse aspects of mRNA lifecycle, including splicing, transport, and stability (Dassi 2017). Diversity in protein architecture and auxiliary domains, as well as a high degree of modularity allow RBPs to impart specific and potent effects on the gene expression of their targets (Janga 2012). For example, the PUF family of proteins in *C. elegans* inhibit translation of their mRNA targets through sequence specific binding of the 3’UTR in order to promote deadenylation or by physically blocking cap recognition by translation initiation factors (reviewed in Wang and Voronina 2020). Other proteins like OMA-1 appear to play a more nuanced role by concomitantly binding the 3’UTRs of mRNAs along with translational repressors like LIN-41 in order to mediate the selective repression-to-activation transition for a subset of mRNAs essential for oogenesis (Tsukamoto *et al.* 2017). Here, RBPs and miRNAs are thought to cooperate extensively, and de-regulation of their activity can precipitate widespread disruption of gene regulatory networks resulting in a variety of cell pathologies and disease states (Tüfekci *et al.* 2013; O’Brien *et al.* 2018).

To effect post-transcriptional regulation of gene expression, RBP and miRNA activity can intersect on multiple levels. On a most basic level, miRNA biogenesis is performed and aided by RBPs (reviewed in Gebert and MacRae 2019). RBPs may directly associate with miRNA-target complexes to modulate the downstream effects on target gene expression (Hammell *et al*., 2009; Schwamborn *et al.* 2009; Wu *et al.* 2017). Coordination between RBPs and miRNAs can also be indirect, with individual factors affecting the target mRNA in distinct ways, ultimately resulting in a unique combinatorial gene regulatory outcome.

Among RBPs identified as modulators of miRNA activity were three proteins that share a conserved RNA-binding K-homology (KH) domain (Akay *et al.* 2013; Zabinsky *et al.* 2017; Li *et al.* 2019). KH domain was first described in human hnRNP K (Siomi *et al.* 1993, reviewed in Geuens *et al.* 2016) and is present alone or in tandem in a large group of RBPs associated with transcription or translation regulation (Nicastro *et al.* 2015; Dominguez *et al.* 2018). The type I KH domain, found in eukaryotes, is approximately 70 amino acids and is characterized by three anti-parallel beta sheets abutted by three alpha helices; it includes the GXXG loop, which is thought to be responsible for nucleic acid binding (Grishin 2001; Valverde *et al.* 2008). We recently showed that HRPK-1, a KH domain-containing protein, physically and functionally interacts with miRNA complexes to modulate gene expression during *C. elegans* development (Li *et al.* 2019). Similarly, the KH domain protein VGLN-1 genetically interacts with a diverse set of miRNAs involved in early embryonic and larval development (Zabinsky *et al.* 2017). VGLN-1 binds mRNAs rich with miRNA binding sites in their 3’UTR (Zabinsky *et al.* 2017) and may serve as a platform, bridging interactions between multiple miRNAs, mRNAs, and proteins to regulate gene expression (Zabinsky *et al.* 2017). GLD-1, an RNA-binding protein and a well-characterized translational repressor that regulates germline development (Marin and Evans 2003), has been shown to genetically interact with multiple miRNAs (Akay *et al.* 2013). GLD-1 contains a single KH domain, functionally interacts with miRNA modulators, *nhl-2* and *vig-1*, and physically interacts with ALG-1, CGH-1, and PAB-1, proteins that are key for miRNA gene regulatory activity (Akay *et al.* 2013). Collectively, these findings suggest that RBPs that harbor KH domain(s) may be functionally important for miRNA-dependent gene regulation.

To determine the extent of functional coordination between the KH domain-containing proteins and miRNAs, we knocked down 24 additional predicted *C. elegans* KH domain genes in sensitized miRNA genetic backgrounds. Strikingly, knock down of 19 KH domain genes resulted in a modulation of a phenotype associated with a partial loss of miRNA activity. We found that several genes, including the predicted splicing factors *sfa-1* and *asd-2* genetically interacted with multiple miRNAs families, suggesting that splicing events may influence miRNA gene regulatory activity. Other genes, such as *Y69A2AR.32*, showed miRNA family specificity. Knockdown of most KH domain genes resulted in enhancement of miRNA reduction-of-function phenotypes, suggesting a normally positive functional interaction between KH domain RBPs and miRNAs. However, knockdown of several genes resulted in mild to strong suppression of defects observed in an Argonaute *alg-1* antimorphic mutant, suggesting that some of these factors normally act antagonistically to miRNAs. Overall, this work provides a comprehensive examination of the possible functional interactions between miRNAs and KH domain RBPs in *C. elegans*, presents a phylogenetic and a domain analysis of *C. elegans* KH domain-containing proteins, and suggests that these RBPs may directly or indirectly coordinate with miRNA pathways to regulate gene expression.

## RESULTS

### Multiple KH domain genes genetically interact with *lsy-6* miRNA in ASEL neuronal cell fate specification

The *lsy-6* miRNA controls cell fate specification of the ASEL/ASER sensory neuron pair. *lsy-6* normally represses expression of *cog-1* in the ASEL neuron, ultimately resulting in an ASEL specific gene expression pattern (Johnston and Hobert 2003) (Figure 1A). Loss of *lsy-6* activity results in an inappropriate cell fate switch of the ASEL neuron to the ASER cell fate (Johnston and Hobert 2003). The *lsy-6(ot150)* reduction-of-function mutation causes a low penetrance phenotype, with ∼15% of *lsy-6(ot150)* animals displaying an ASEL cell fate defective phenotype. This cell fate defect can be observed by the loss of the *Plim-6::gfp* expression within the ASEL neuron (Figure 1A, B).

**Figure 1.**
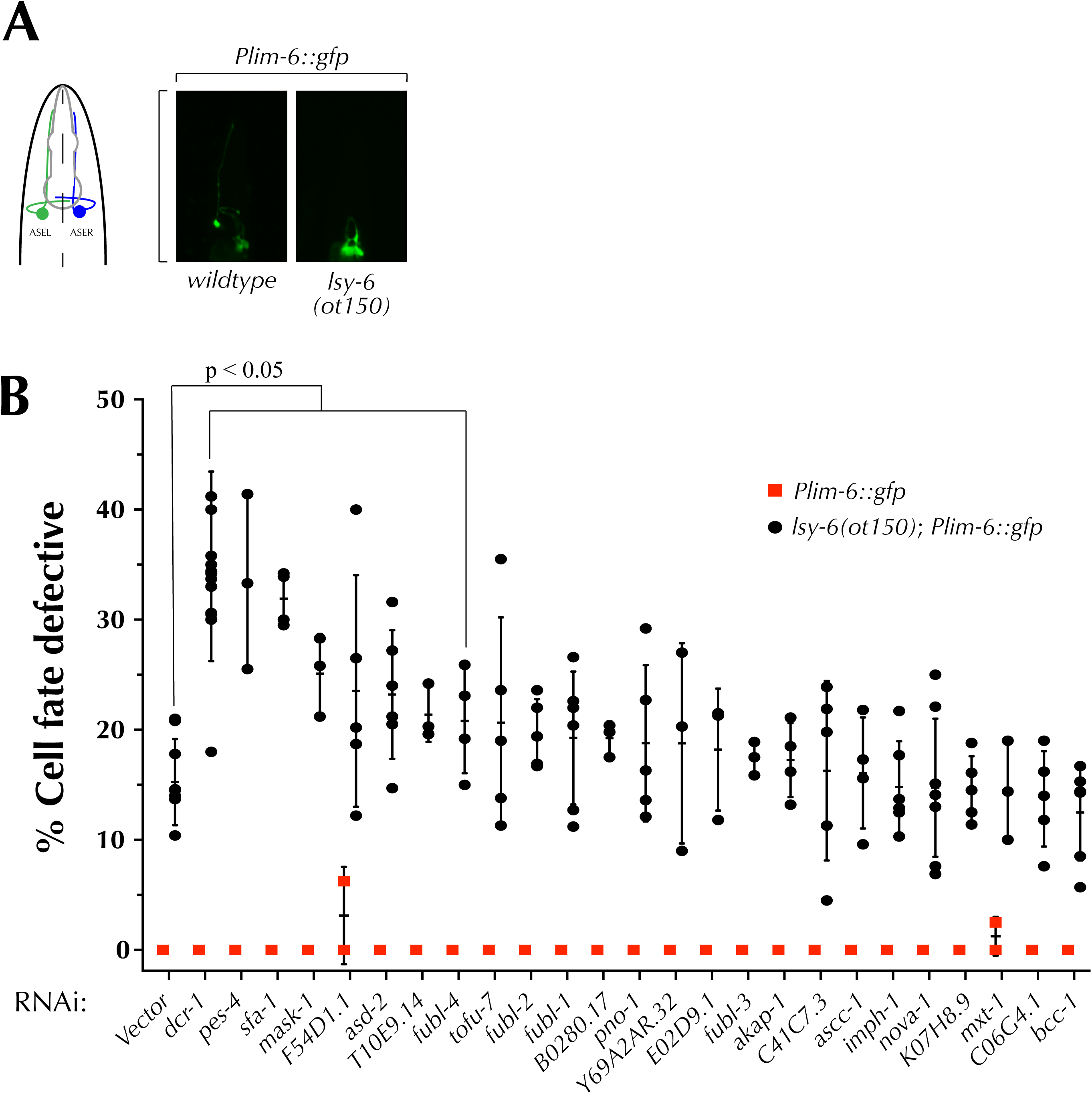
Knockdown of several KH domain genes enhances the cell defective phenotype of *lsy-6(ot150)* mutants. (**A**) *lsy-6* miRNA directs the ASEL cell fate specification, with ASEL cell fate marked by the *lim-6::gfp* reporter. *lsy-6(ot150)* mutations results in partially penetrant loss of *lim-6::gfp* expression in ASEL cells. (**B**) RNAi-mediated knockdown of seven KH domain genes significantly enhances the cell fate defective phenotype of *lsy-6(ot150)* animals. Dots represent experimental replicates. Unpaired t-test was used to compare gene RNAi to vector control within individual genotypes.

To identify whether KH-domain genes play a role in *lsy-6*-dependent neuronal cell fate specification, we knocked down 24 KH domain genes in the *lsy-6(ot150)* mutant background (Figure 1B) and assayed the penetrance of the ASEL cell fate defect. Knockdown of seven of the KH domain genes, *pes-4, sfa-1, mask-1, F54D1.1, asd-2, T10E9.14, and fubl-4*, significantly enhanced the *lsy-6(ot150)* cell fate defective phenotype (Figure 1B, Table 1). RNAi-mediated knockdown of the same genes did not result in a phenotype in the absence of the *lsy-6(ot150)* allele, with the exception of *F54D1.1* and *mxt-1*, whose depletion resulted in occasional loss of *Plim-6::gfp* expression in ASEL (Figure 1B). Furthermore, RNAi of additional genes, such as *tofu-*7 and *pno-1*, resulted in a mild but overall not statistically significant enhancement of the *lsy-6(ot150)* phenotype. These genetic interactions may nevertheless be biologically relevant.

**Table 1.** KH domain genes functionally interact with miRNA sensitized mutants.

| Gene or allele | *lsy-6(ot150)* | *lsy-6* | *mir-48 mir-241(nDf51)* |  | *let-7(n2853)* | *alg-1(ma202)* |
| --- | --- | --- | --- | --- | --- | --- |
| Assay | Cell fate<sup>a</sup><br>defective | Uterine<sup>b</sup><br>*cog-1::gfp* | Abnormal<sup>c</sup><br>*col-19::GFP* | Seam<sup>d</sup><br>Cell Number | Bursting<sup>e</sup> | Wildtype<sup>f</sup><br>*col-19::gfp* |
| RNAi |  |  |  |  |  |  |
| *Empty vector* | 15.2 ± 3.9 (n=688) | 48.8 ± 1.5 (n=68) | 11.2 ± 7.9 (n=149) | 13.4 ± 1.4 | 13.7 ± 6.9 (n=309) | 0 (n=33) |
| *dcr-1* | 34.9 ± 8.6 (n=935) | 60.2 ± 10.8 (n=106) | 79.4 ± 15.7 (n=186) | 14.6 ± 1.9 | 33.4 ± 4.8 (n=230) | n.d. |
| *fubl-3* | 17.4 ± 1.5 (n=192) | n.d.<sup>g</sup> | 50.4 ± 19.2 (n=44)<sup>h</sup> | 14.2 ± 1.7 | 31.7 ± 8.7 (n=145) | 0 (n=31) |
| *fubl-4* | 20.8 ± 4.7 (n=415) | 46.6 ± 17.7 (n=63) | 49.7 ± 7.5 (n=40) | 14.1 ± 2.1 | 14.5 ± 10.6 (n=175) | 0 (n=32) |
| *fubl-1* | 19.3 ± 6.0 (n=570) | n.d. | 37.2 ± 29.5 (n=83) | 13.3 ± 1.4 | 24.2 ± 6.2 (n=197) | 2.7 ± 3.8 (n=28) |
| *fubl-2* | 19.7 ± 3.1 (n=485) | n.d. | 22.3 ± 19.7 (n=45) | 13.2 ± 1.5 | 12.2 ± 2.5 (n=200) | 2.3 ± 3.2 (n=30) |
| *imph-1* | 14.8 ± 4.2 (n=634) | n.d. | 6.2 ± 5.4 (n=45) | 13.5 ± 1.5 | 9.5 ± 1.8 (n=190) | 7.3 ± 7.2 (n=40) |
| *pes-4* | 33.4 ± 8.0 (n=113) | 66.2 ± 7.4 (n=32) | 34.0 ± 12.2 (n=38) | 12.7 ± 1.1 | 10.6 ± 2.1 (n=349) | 0 (n=20) |
| *T10E9.14* | 21.4 ± 2.5 (n=175) | 63.9 ± 3.9 (n=36) | 51.9 ± 18.9 (n=36) | 14.1 ± 1.9 | 15.9 ± 8.3 (n=160) | 0 (n=35) |
| *nova-1* | 14.7 ± 6.3 (n=965) | n.d. | 6.7 ± 9.4 (n=25) | 14.0 ± 1.6 | 11.5 ± 4.8 (n=195) | 3.0 ± 4.2 (n=33) |
| *mxt-1* | 14.5 ± 4.5 (n=249) | n.d. | 26.6 ± 14.7 (n=64) | 14.2 ± 1.9 | 28.4 ± 11.9 (n=192) | 0 (n=30) |
| *ascc-1* | 16.1 ± 5.1 (n=257) | n.d. | 18.1 ± 13.5 (n=61) | 13.2 ± 1.3 | 15.2 ± 7.1 (n=159) | 4.2 ± 5.9 (n=18) |
| *akap-1* | 17.3 ± 3.4 (n=402) | n.d. | 35.8 ± 2.1 (n=57) | 14.1 ± 1.5 | 13.2 ± 9.4 (n=198) | 7.2 ± 10.1 (n=42) |
| *tofu-7* | 20.6 ± 9.6 (n=532) | 71.6 ± 1.1 (n=50) | 37.2 ± 18.2 (n=53) | 14.2 ± 2.1 | 17.5 ± 6.5 (n=148) | 0 (n=34) |
| *C06G4.1* | 13.7 ± 4.3 (n=395) | n.d. | 13.9 ± 4.0 (n=33) | 13.4 ± 1.6 | 10.9 ± 6.9 (n=157) | 2.3 ± 3.2 (n=30) |
| *E02D9.1* | 18.2 ± 5.5 (n=160) | n.d. | 3.9 ± 6.8 (n=60) | 13.2 ± 1.2 | 7.2 ± 5.4 (n=190) | 58.0 ± 19.0 (n=26) |
| *sfa-1* | 32.0 ± 2.5 (n=152) | 73.0 ± 9.0 (n=41) | 43.9 ± 27.3 (n=78) | 14.4 ± 1.8 | 25.3 ± 19.2 (n=210) | 7.2 ± 6.2 (n=31) |
| *asd-2* | 23.2 ± 5.8 (n=375) | 65.4 ± 1.7 (n=49) | 51.8 ± 23.9 (n=63) | 13.9 ± 1.9 | 30.6 ± 19.8 (n=188) | 0 (n=42) |
| *B0280.17* | 19.3 ± 1.5 (n=220) | n.d. | 29.9 ± 19.8 (n=52) | 13.5 ± 1.5 | 31.5 ± 9.2 (n=98) | 0 (n=30) |
| *F54D1.1* | 23.5 ± 10.5 (n=659) | 58.3 ± 25.1 (n=61) | 24.9 ± 7.7 (n=53) | 13.9 ± 1.4 | 15.7 ± 7.0 (n=206) | 0 (n=29) |
| *K07H8.9* | 14.7 ± 2.9 (n=340) | n.d. | 10.8 ± 14.3 (n=62) | 13.7 ± 1.4 | 9.7 ± 5.4 (n=197) | 3.0 ± 4.1 (n=33) |
| *Y69A2AR.32* | 18.8 ± 9.1 (n=163) | n.d. | 40.8 ± 4.6 (n=49) | 13.5 ± 1.4 | 12.1 ± 3.0 (n=145) | 6.0 ± 1.6 (n=35) |
| *bcc-1* | 12.5 ± 4.4 (n=745) | n.d. | 20.1 ± 12.5 (n=85) | 14.0 ± 1.9 | 31.0 ± 12.5 (n=148) | 0 (n=26) |
| *C41G7.3* | 16.3 ± 8.2 (n=364) | n.d. | 45.5 ± 15.4 (n=68) | 15.0 ± 2.3 | 13.3 ± 9.2 (n=199) | 1.4 ± 2.5 (n=38) |
| *pno-1* | 18.8 ± 7.1 (n=523) | n.d. | 40.2 ± 12.0 (n=85) | 13.3 ± 1.4 | 14.0 ± 6.8 (n=200) | 0 (n=25) |
| *mask-1* | 25.1 ± 3.6 (n=193) | 60.6 ± 5.8 (n=40) | 50.7 ± 18.6 (n=37) | 14.7 ± 2.3 | 12.9 ± 3.0 (n=149) | 0 (n=25) |

<sup>a</sup> *Plim-6::gfp* was scored in L4-Adult worms; n > 160 (range 160-965)
<sup>b</sup> Top enhancers of *lsy-6* activity were scored for uterine *cog-1::gfp* expression in L4 worms; n > 36 (range 36-106)
<sup>c</sup> *col-19::gfp* was scored in young adult worms; n > 25 (range 25-186)
<sup>d</sup> Seam cell number was scored at the same time as *col-19::GFP* expression in young adults; n > 25 (range 25-186)
<sup>e</sup> Synchronized L1 worms were reared at 15° C; vulval bursting scored in day 1 adults; n > 98 (range 98-308)
<sup>f</sup> *col-19::gfp* was scored in young adult worms; n > 18 (range 18-40)
<sup>g</sup> n.d. - not determined
<sup>h</sup> values showing statistical significance are highlighted in red

### KH domain genes coordinate with *lsy-6* to regulate the expression of *cog-1*

Next, we wanted to determine whether the genes that genetically interacted with *lsy-6(ot150)* were also able to regulate a *lsy-6* target, *cog-1* (Johnston and Hobert 2003). While *lsy-6* expression is normally restricted to neuronal tissues, its endogenous target *cog-1* is more broadly expressed (Palmer *et al.* 2002). Expression of *lsy-6* from the *cog-1* promoter represses the *cog-1::gfp* reporter in the uterine and vulval cells (Johnston and Hobert 2003) (Figure 2A). Therefore, we can utilize the *lsy-6*-mediated repression of *cog-1* to assay the effects of knocking down potential modulators of *lsy-6* activity. Indeed, RNAi of six out of eight genes that showed genetic interactions with *lsy-6* (Figure 1B), *sfa-1, tofu-7, pes-4, asd-2, T10E9.14*, and *mask-1* resulted in a significant de-repression of *cog-1::gfp* expression in uterine cells (Figure 2B, Table 1), suggesting that these genes may coordinate with *lsy-6* in repressing *cog-1*. However, knockdown of *tofu-7* repressed *cog-1* expression in the uterine cells in the absence of *Pcog-1::lsy-6* (Figure 2B), suggesting that *tofu-7* may regulate *cog-1* independently of *lsy-6.* Knockdown of *asd-2*, and *F54D1.1* produced similar, albeit not statistically significant, effects on *cog-1::gfp* expression independent of *lsy-6* (Figure 2B). In fact, *tofu-7, asd-2*, and *F54D1.*1 may have a more complex functional relationship, perhaps regulating *cog-1* through multiple genetic pathways, one of which involves the *lsy-6* miRNA. Here, in the absence of *lsy-6, tofu-7, asd-2*, and *F54D1.1* may act to promote *cog-1::gfp* expression, while the addition of *lsy-6* changes the functional relationship from positive to repressive or may de-regulate target gene expression in either direction (Figure 2B).

**Figure 2.**
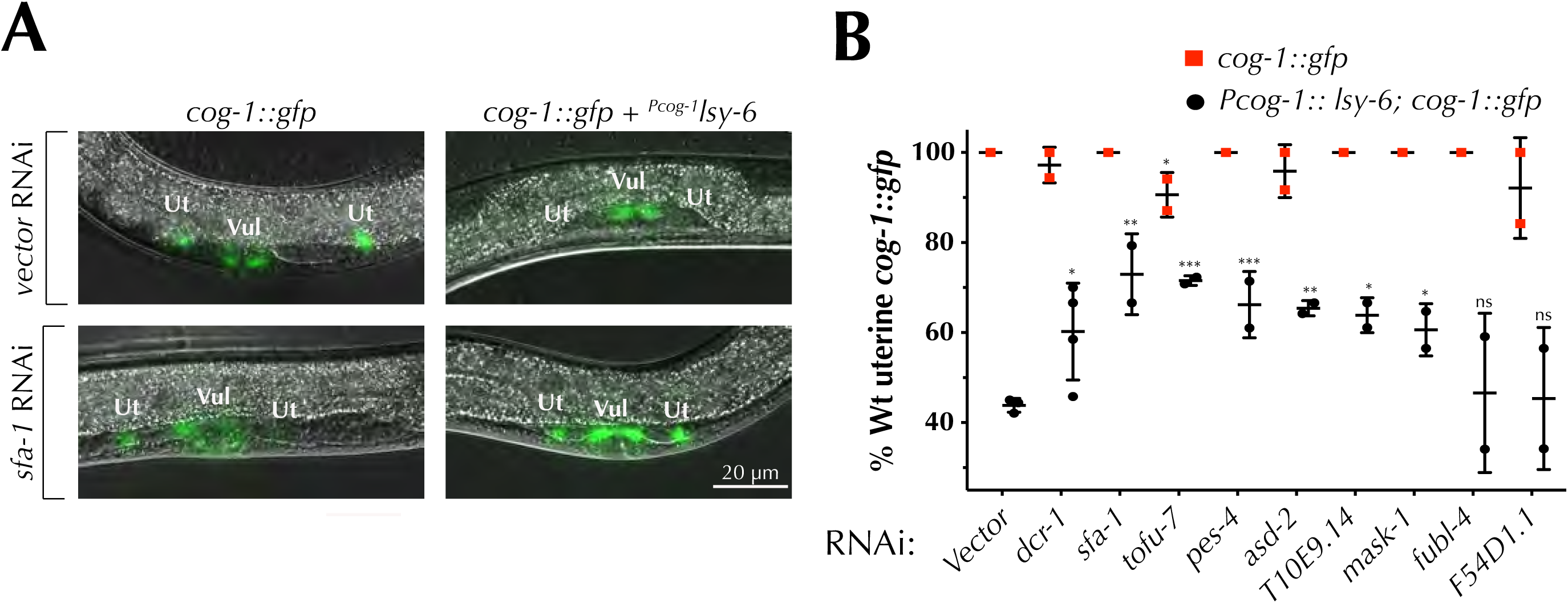
Several KH domain genes coordinate with *lsy-6* to regulate *cog-1::gfp* expression in uterine cells. RNAi of several KH domain genes, including *sfa-1* (**A**) alleviates the *lsy-6*-mediated repression of *cog-1*::*gfp* in uterine cells (**B**). Unpaired t-test was used to compare gene RNAi to vector control within each individual genotype. Dots represent experimental replicates.

### KH domains proteins coordinate with *let-7* family of miRNAs

To determine whether KH domain-containing proteins might coordinate with additional miRNAs beyond *lsy-6*, we looked for a genetic interaction between the KH domain genes and the *let-7*-family of miRNAs. The *let-7* miRNA family, as part of a complex genetic network, regulates division patterns and terminal cell differentiation of seam cells during *C. elegans* larval development (Reinhart *et al.* 2000; Slack *et al.* 2000; Abbott *et al.* 2005). Three members of the *let-7* family, *mir-48, mir-241*, and *mir-84* act redundantly to control seam cell divisions by inhibiting the proliferative divisions of the L2 stage and promoting the self-renewing seam cell divisions of the L3 stage (Abbott *et al.* 2005). Loss of *mir-48, mir-241*, and *mir-84* leads to a highly penetrant reiteration of the proliferative L2 seam cell division leading to increased seam cell number, delayed alae formation, and delayed expression of the adult specific reporter, *col-19::gfp* (Abbott *et al.* 2005). Deletion of *mir-48* and *mir-241*, which leaves *mir-84* intact, results in milder heterochronic phenotypes including increased seam cell number and retarded alae formation (Abbott *et al.* 2005).

We performed RNAi for the 24 KH domain genes in the *mir-48 mir-241(nDf51)* mutant background and assayed both *col-19::gfp* expression and seam cell number in young adult animals (Figure 3A, B). RNAi of 15 KH domain genes significantly enhanced the retarded *col-19::gfp* expression observed in *mir-48 mir-241(nDf51)* animals (Figure 3B, Table 1). RNAi of these genes did not produce a phenotype in the absence of the *mir-48 mir-241* deletion (Figure 3B), with the exception of *B0280.17* RNAi, which exhibited a very mild defect in hypodermal *col-19::gfp* expression. In addition, *F54D1.1* RNAi produced a mildly penetrant abnormal *col-19::gfp* expression, but did not enhance the *mir-48 mir-241* phenotype to a statistically significant level (Figure 3B). RNAi of 13 KH domain genes produced a significant increase in the average number of seam cells in the *mir-48 mir-241(nDf51)* mutants compared to the empty vector control (Figure 3C, Table 1). Overall, depletion of ten genes enhanced the delayed hypodermal *col-19::gfp* expression and increased the seam cell number of *mir-48 mir-241(nDf51)* mutants (Figure 3B, C, Table 1). Together these data suggest that a subset of KH domain genes may coordinate, directly or indirectly, with the *let-7 family* miRNAs to regulate their target gene expression.

**Figure 3.**
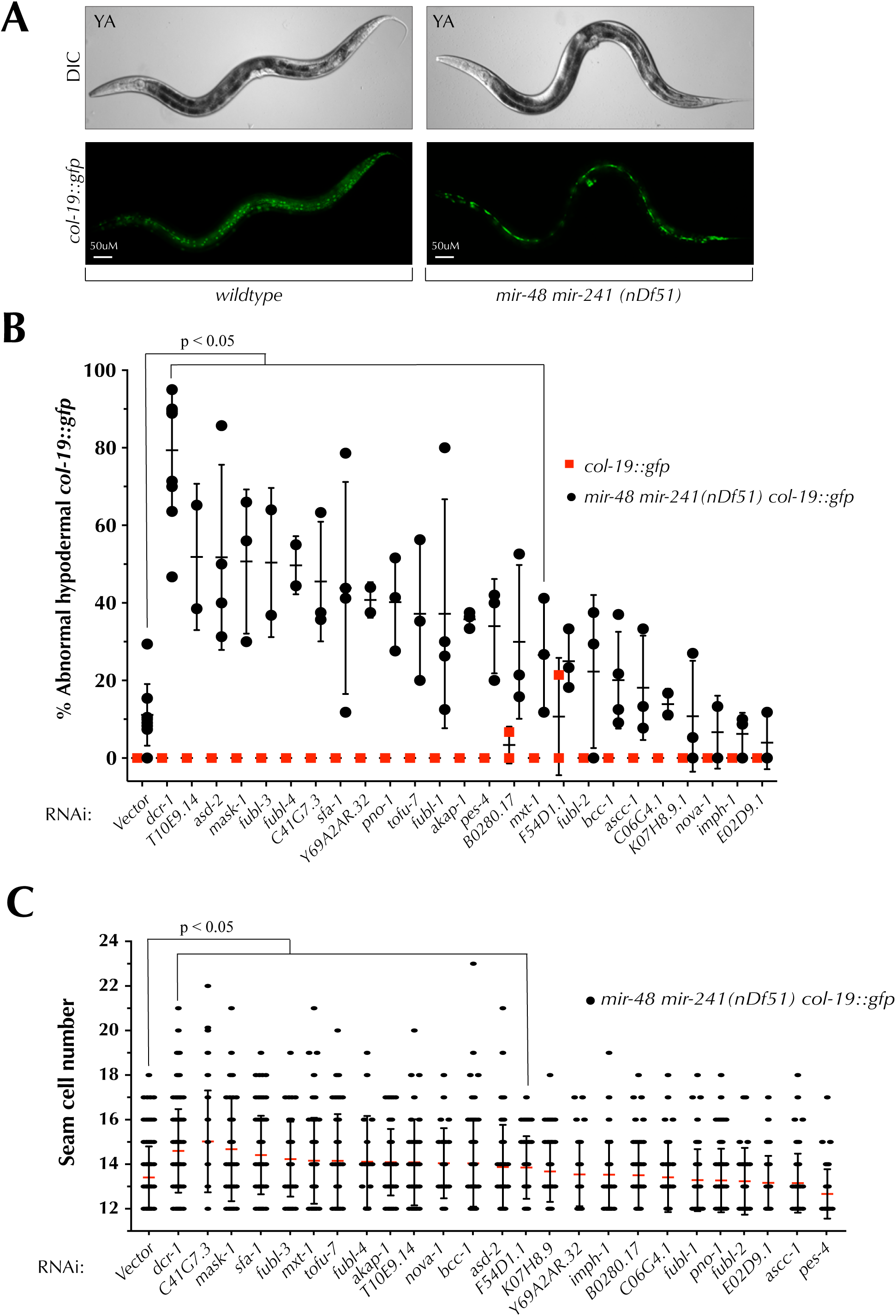
RNAi of multiple KH domain genes enhances the *mir-48 mir-241* heterochronic phenotype. (**A**) Loss of *mir-48 mir-241* results in delayed hypodermal expression of the adult specific marker *col-19::gfp*. (**B**) When compared to vector RNAi knockdown of 15 KH domain genes by RNAi enhances the retarded hypodermal *col-19::gfp* expression of *mir-48 mir-241(nDf51)* animals. Dots represent experimental replicates. (**C**) RNAi of some KH domain genes increases the seam cell numbers of *mir-48 mir-241(nDf51)* young adults when compared to vector RNAi. Each dot represents the seam cell number of an individual worm after RNAi. Statistical significance determined by unpaired t-test.

To assess the functional relevance of KH domain genes to miRNA activity later in development, we asked whether reducing KH domain gene function impacts activity of *let-7* itself. *let-7* governs the terminal seam cell differentiation during the transition from L4 to adulthood (Reinhart *et al.* 2000). Compromising *let-7* miRNA activity produces a heterochronic phenotype, which, among other defects, includes vulval rupture during the L4-adult molt (Reinhart *et al.* 2000). *let-7(n2853)*, a temperature sensitive reduction-of-function mutation, causes a mildly penetrant vulval rupture phenotype at 15°C (Figure 4A) (Reinhart *et al.* 2000). RNAi of seven KH domain genes led to significant enhancement of the vuval bursting phenotype (Figure 4B) suggesting these genes may coordinate with *let-7* miRNA in a way that normally promotes its activity. RNAi of two genes (*Y69A2AR.32* and *K07H8.9)* resulted in vulval bursting in a small percentage of wildtype animals, suggesting they may regulate vulval development independently of *let-7*.

**Figure 4.**
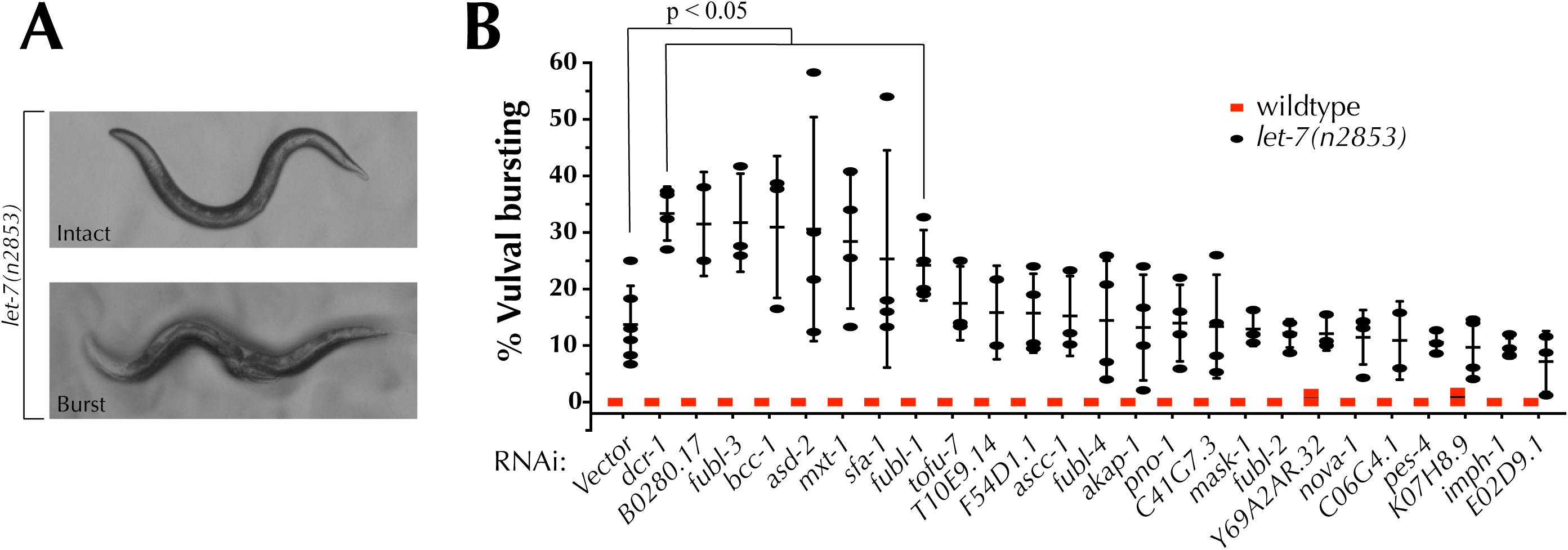
Several KH domain genes interact genetically with the *let-7* miRNA. (**A, B**) *let-5(n2853)* worms display a partially penetrant vulval bursting phenotype at permissive temperature (15°C). (**B**) RNAi knockdown of seven KH domain genes significantly enhances the vulval bursting phenotype of *let-7(n2853)* worms. Statistical significance determined by unpaired t-test. Dots represent experimental replicates.

### Knockdown of several KH domain genes suppresses compromised miRISC activity

ALG-1 is one of two *C. elegans* Argonautes (ALG-1 and ALG-2) that primarily associate with miRNAs and are central for miRNA biogenesis and activity (Grishok *et al.* 2005). Mutations abolishing ALG-1 activity result in moderate developmental defects, while abolishing both *alg-1* and *alg-2* activity results in early lethality (Grishok *et al.* 2005; Vasquez-Rifo *et al.* 2012). In addition, antimorphic mutations in *alg-1*, such as *alg-1(ma202)*, result in more pronounced defects in miRNA activity, likely due to sequestration of miRNA pathway components away from ALG-2 (Zinovyeva *et al.* 2014). Specifically, *alg-1(ma202)* animals display severe heterochronic defects (Zinovyeva *et al.* 2014), with 100% of *alg-1(ma202)* young adult animals failing to appropriately express adult cell marker *col-19::gfp* (Zinovyeva *et al.* 2014) (Figure 5A,B).

**Figure 5.**
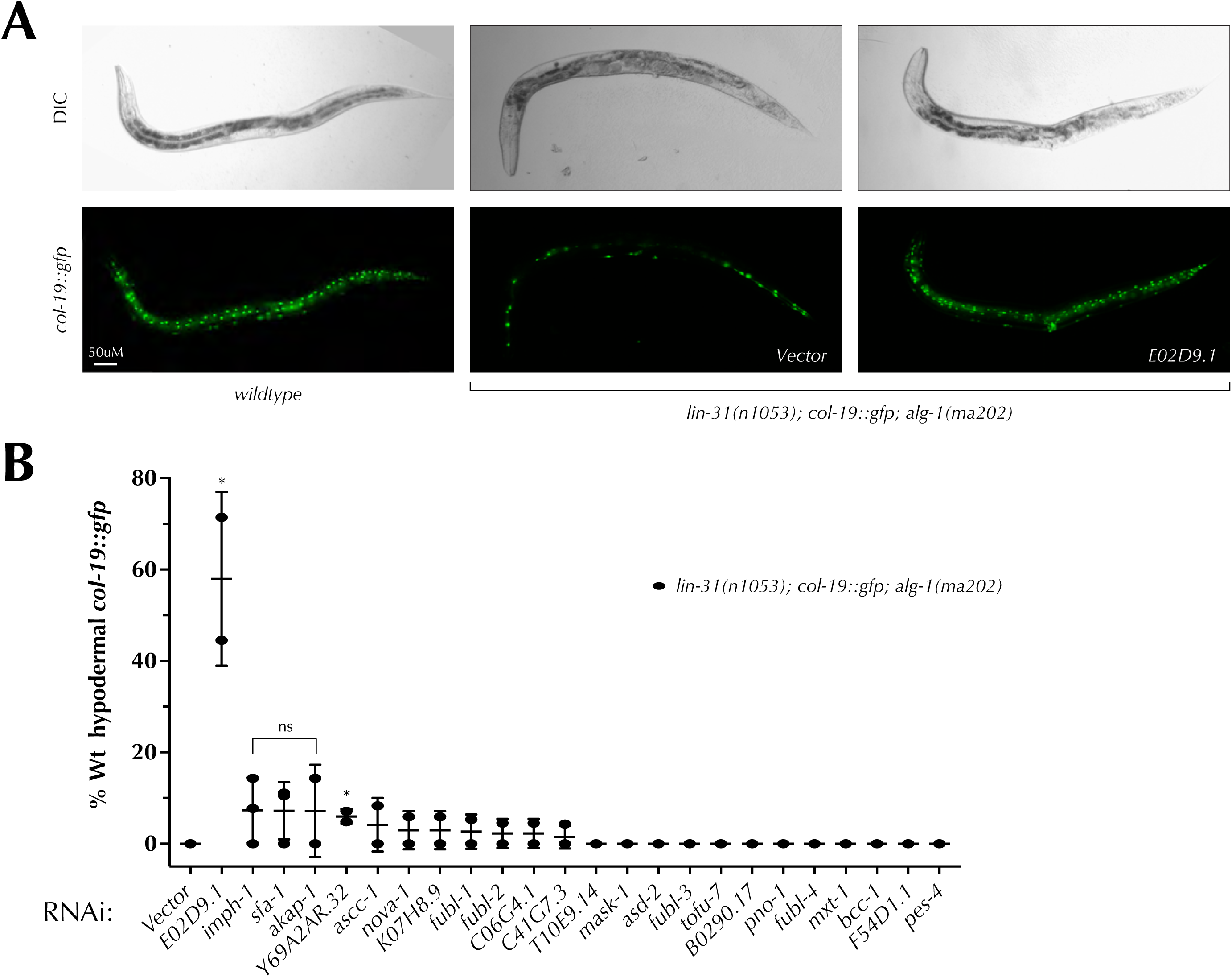
Two KH domain containing genes interact genetically with ALG-1(*ma202*). (**A**) *alg-1(ma202)* young adults lack hypodermal *col-19::gfp* expression. The *alg-1(ma202)* mutation is present in a *lin-31(n1053)* background to suppress bursting via non-heterochronic mechanisms. RNAi of *E02D9.1* restores *col-19::gfp* expression in young adults (**A**,**B**). (**B**) RNAi of several genes suppresses the retarded *col-19::gfp* expression phenotype of *alg-1(ma202)* mutants. Statistical significance determined by unpaired t-test. Dots represent experimental replicates.

To test whether KH domain genes genetically interact with the *alg-1(ma202)* allele, we performed RNAi knockdown in the *alg-1(ma202)* background and assessed *col-19::gfp* expression in hypodermal cells of young adult animals. We used this background to screen for genes that may normally negatively interact with the miRNA pathways and therefore suppress the *alg-1(ma202)* phenotype when knocked down. Interestingly, RNAi of two genes, *E02D9.1* and *Y69A2AR.32* significantly suppressed the abnormal hypodermal *col-19::gfp* expression in *alg-1(ma202)* young adults (Figure 5B, Table 1). Especially striking is the *E02D9.1* RNAi suppression, which results in ∼ 60% of *alg-1(ma202)* animals exhibiting wild type hypodermal *col-19::gfp* expression as young adults (Figure 5A,B). Although not statistically significant, possibly due to the variation in RNAi efficiency, RNAi of several other genes (*imph-1, sfa-1*, and *akap-1)* restored wild type *col-19::gfp* expression in *alg-1(ma202)* young adults, something that is never observed in *alg-1(ma202)* mutants alone (Figure 5B). As *alg-1(ma202)* suppressors, these KH domain genes may act in a manner that opposes normal miRNA activity, with their depletion perhaps resulting in decreased miRNA target gene expression.

### KH domain containing RBPs play a role in early development

To determine whether KH domain genes have a general effect on *C. elegans* development, we assayed the brood size and embryonic lethality of animals with reduced KH domain gene function. Knockdown of 10 genes (*fubl-3, fubl-4, pes-4, akap-1, tofu-7, E02D9.1, sfa-1, Y69A2AR.32, bcc-1*, and *mask-1*) resulted in significant reduction in brood size (Figure 6A, Table 2), while knockdown of 8 genes (*pes-4, T10E9.14, akap-1, tofu-7, sfa-1, asd-2, K07H8.9*, and *bcc-1*) causes significant embryonic lethality (Figure 6B, Table 2). Depletion of *sfa-1, pes-4, akap-1, tofu-7*, and *bcc-1* had significant effects on both brood size and embryonic lethality suggesting that these genes play fundamental roles in *C. elegans* development (Figure 6 A, B, Table 2). Several additional genes disrupted early development, albeit to a degree that was not statistically significantly by our analysis (Figure 6, Table 2). These observations are consistent with previously reported roles for *akap-1, E02D9.1, sfa-1, asd-2, K07H8.9*, and *bcc-1* in early *C. elegans* development (Kamath *et al.* 2003; Sönnichsen *et al.* 2005; Ohno *et al.* 2008; Ma and Horvitz 2009; Kapelle and Reinke 2011) and highlight additional genes as important for *C. elegans* fecundity and embryonic development.

**Table 2.** Knockdown of KH domain genes affects embryonic lethality and brood size.

| RNAi \ Assay | Embryonic lethality<sup>a</sup> | Brood size<sup>b</sup> |
| --- | --- | --- |
| *Empty vector* | 1.6 ± 1.3 | 338.3 ± 36.5 |
| *dcr-1* | 6.9 ± 4.6 | 280.0 ± 36.6 |
| *fubl-3* | 5.0 ± 3.3 | 267.7 ± 43.9 |
| *fubl-4* | 4.8 ± 5.4 | 237.7 ± 61.4 |
| *fubl-1* | 2.1 ± 2.0 | 324.0 ± 49.1 |
| *fubl-2* | 5.8 ± 3.3 | 311.4 ± 44.1 |
| *imph-1* | 6.2 ± 5.4 | 284.1 ± 10.3 |
| *pes-4* | 22.8 ± 30.9 | 7.2 ± 9.9 |
| *T10E9.14* | 11.5 ± 5.4 | 271.3 ± 67.9 |
| *nova-1* | 6.2 ± 4.3 | 306.3 ± 46.8 |
| *mxt-1* | 4.4 ± 4.1 | 322.3 ± 41.6 |
| *ascc-1* | 3.1 ± 2.2 | 292.4 ± 27.2 |
| *akap-1* | 13.6 ± 9.1 | 199.3 ± 66.0 |
| *tofu-7* | 11.5 ± 8.2 | 265.3 ± 100.0 |
| *C06G4.1* | 5.4 ± 6.0 | 313.2 ± 25.3 |
| *E02D9.1* | 5.9 ± 6.0 | 205.6 ± 140.2 |
| *sfa-1* | 37.1 ± 13.6 | 221.8 ± 56.8 |
| *asd-2* | 8.3 ± 4.0 | 321.8 ± 47.9 |
| *B0280.17* | 4.2 ± 2.6 | 275.8 ± 16.8 |
| *F54D1.1* | 3.4 ± 2.7 | 306.8 ± 33.1 |
| *K07H8.9* | 12.2 ± 14.7 | 275.8 ± 64.8 |
| *Y69A2AR.32* | 1.2 ± 1.0 | 257.8 ± 70.2 |
| *bcc-1* | 6.9 ± 2.3 | 253.4 ± 57.1 |
| *C41G7.3* | 2.6 ± 2.3 | 307.4 ± 71.1 |
| *pno-1* | 4.9 ± 2.3 | 294.3 ± 26.5 |
| *mask-1* | 1.7 ± 0.6 | 262.4 ± 33.3 |

<sup>a</sup> All worms were grown at 20° C; dead embryos were counted 1 day after being laid; n = number of broods; n > 4 (range 4-13)
<sup>b</sup> All worms were grown at 20° C; brood size includes dead embryos, arrested larvae and living progeny; n = number of broods; n > 4 (range 4-13)

**Figure 6.**
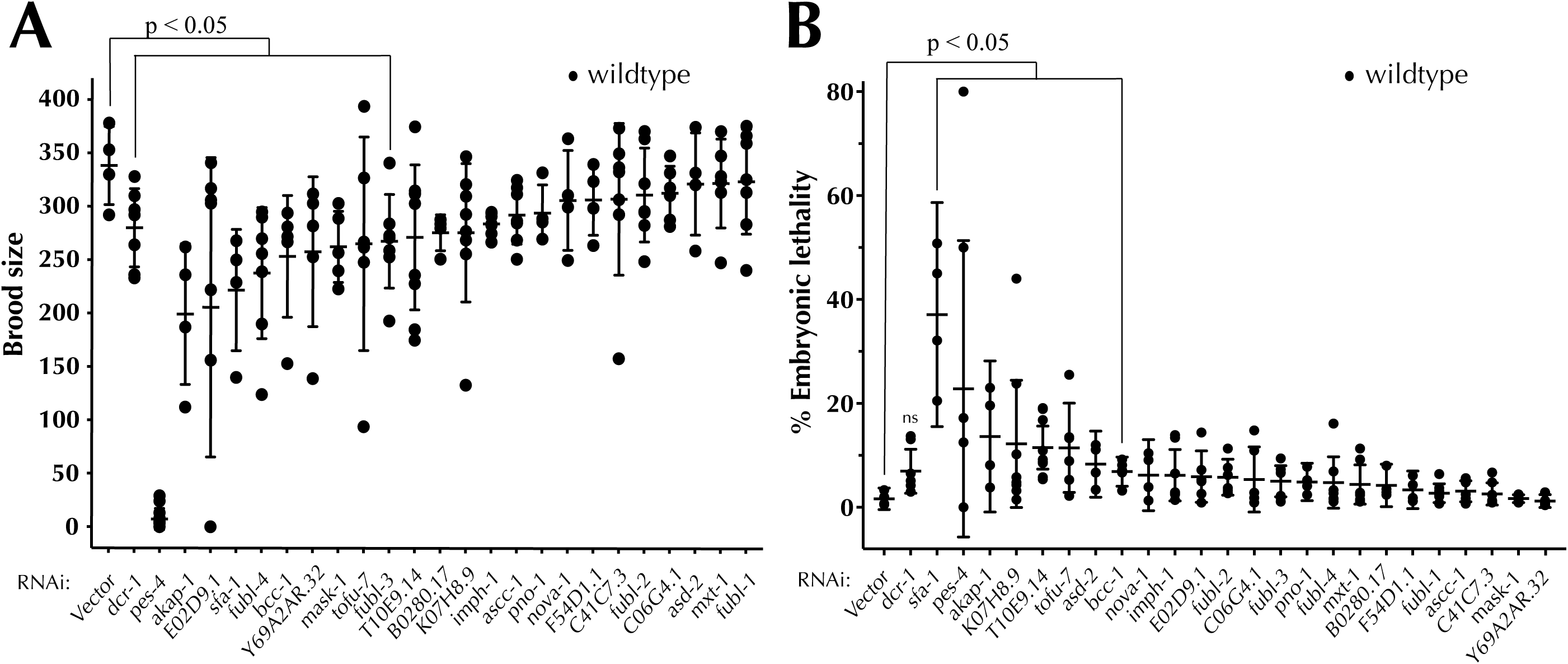
KH domain containing RNA-binding proteins may have essential roles in development. (**A**) RNAi of 10 KH domain genes resulted in significant reductions in brood size. (**B**) Depletion of eight KH domain proteins results in significant embryonic lethality resulted. Statistical significance determined by unpaired t-test.

### Some *C. elegans* KH domain proteins are evolutionary related and have diverse domain architecture

Protein domains are discrete functional and structural segments of a protein. The loss, gain, or structural modification of domains can drive evolution, allowing proteins to lose or acquire new functions over evolutionary time. As domains evolve from ancestral forms, proteins containing the same types of domains may be evolutionary related. To understand the evolutionary relationship between the KH domain-containing proteins and to potentially inform our functional analysis, we performed an alignment of *C. elegans* KH domain protein sequences using the MEGAx alignment program (Kumar *et al.* 2018) and generated a phylogenetic tree (Figure 7). Interestingly, proteins that appear to coordinate with miRNAs are found in almost every clade in our phylogenetic analysis (Figure 7).

**Figure 7.**
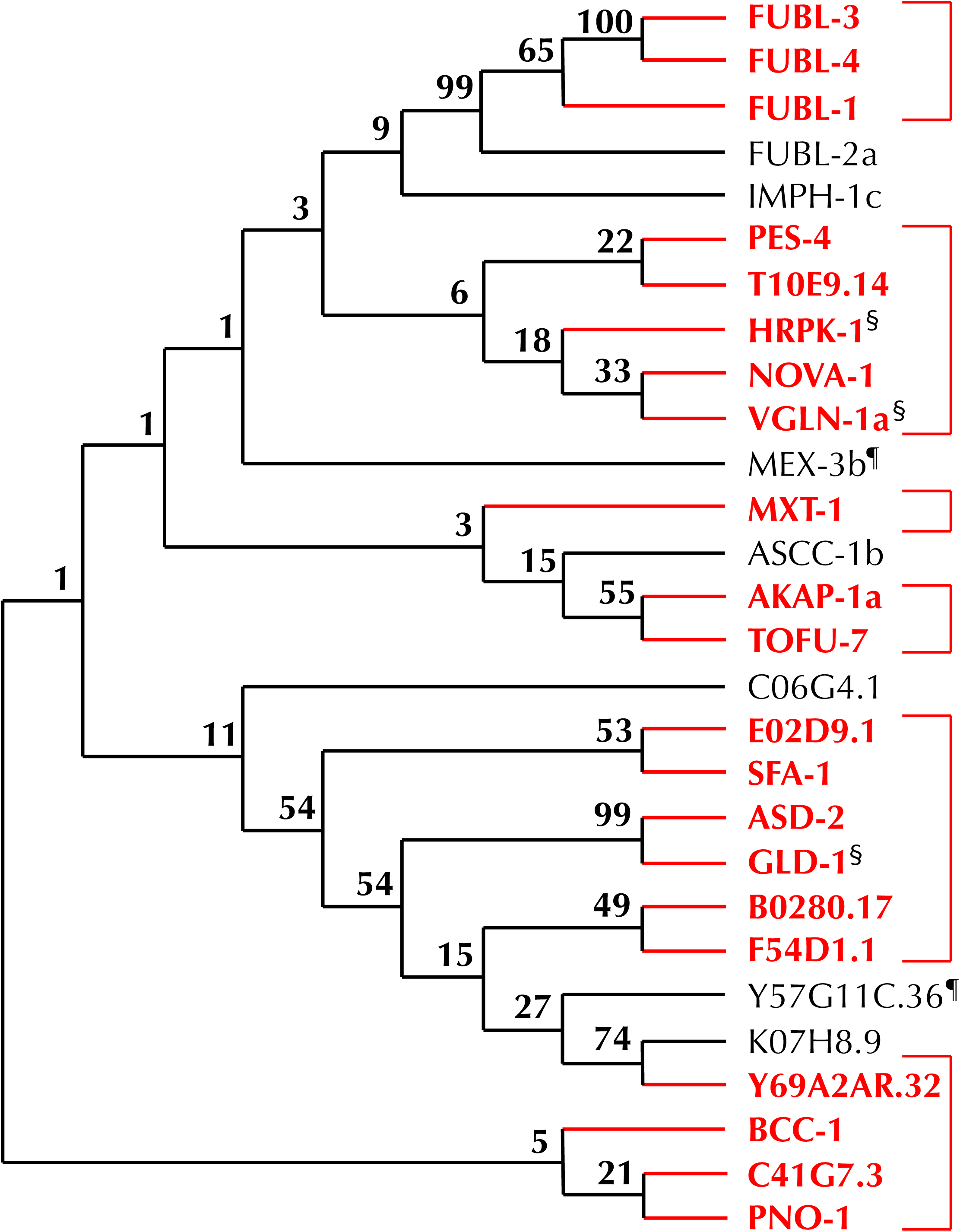
Phylogenetic analysis of KH domain containing RNA-binding proteins. Multiple sequence alignment of 28 KH domain containing RNA-binding proteins was performed and the proteins grouped in clades based on sequence similarity; branches are labeled with confidence value. Clades containing proteins that genetically interact with one or more miRNA sensitized background are bracketed and highlighted in red. A ¶ indicates that functional assays were not performed for a particular gene. A § denotes genes identified as interacting with miRNAs in other publications.

KH domains are thought to mediate numerous interactions, including those between proteins (Valverde *et al.* 2008) and proteins and nucleic acids (Grishin 2001; Valverde *et al.* 2008). Due to the KH domain’s ability to bind RNA, the *C. elegans* KH domain-containing proteins represent a subset of RBPs, but combinatorial domain arrangements can result in extensive functional diversity among them. To determine the diversity of domain structures of KH domain containing proteins, we analyzed their domain architecture using the Simple Modular Architecture Research Tool (SMART) (Letunic and Bork 2017), which identifies known domain sequences. In addition, we utilized the PLACC web-based tool to identify prion-like domains, or unstructured regions (Lancaster *et al.* 2014). Such low complexity regions are thought to have affinity for RNA (Kato *et al.* 2012) and can play a role in phase-phase separation that is important for forming and reforming of ribonucleoprotein (RNP) bodies (Shin and Brangwynne 2017). We found that KH domain-containing proteins harbor a diverse set of domains, with prion-like domains present in 17/29 of KH domain proteins (Figure 8). Unsurprisingly, many proteins of the same clade shared additional domains (Figure 7, Figure 8). These analyses may in the future may help inform the mechanisms by which these proteins coordinate with miRNA-mediated regulation of gene expression.

**Figure 8.**
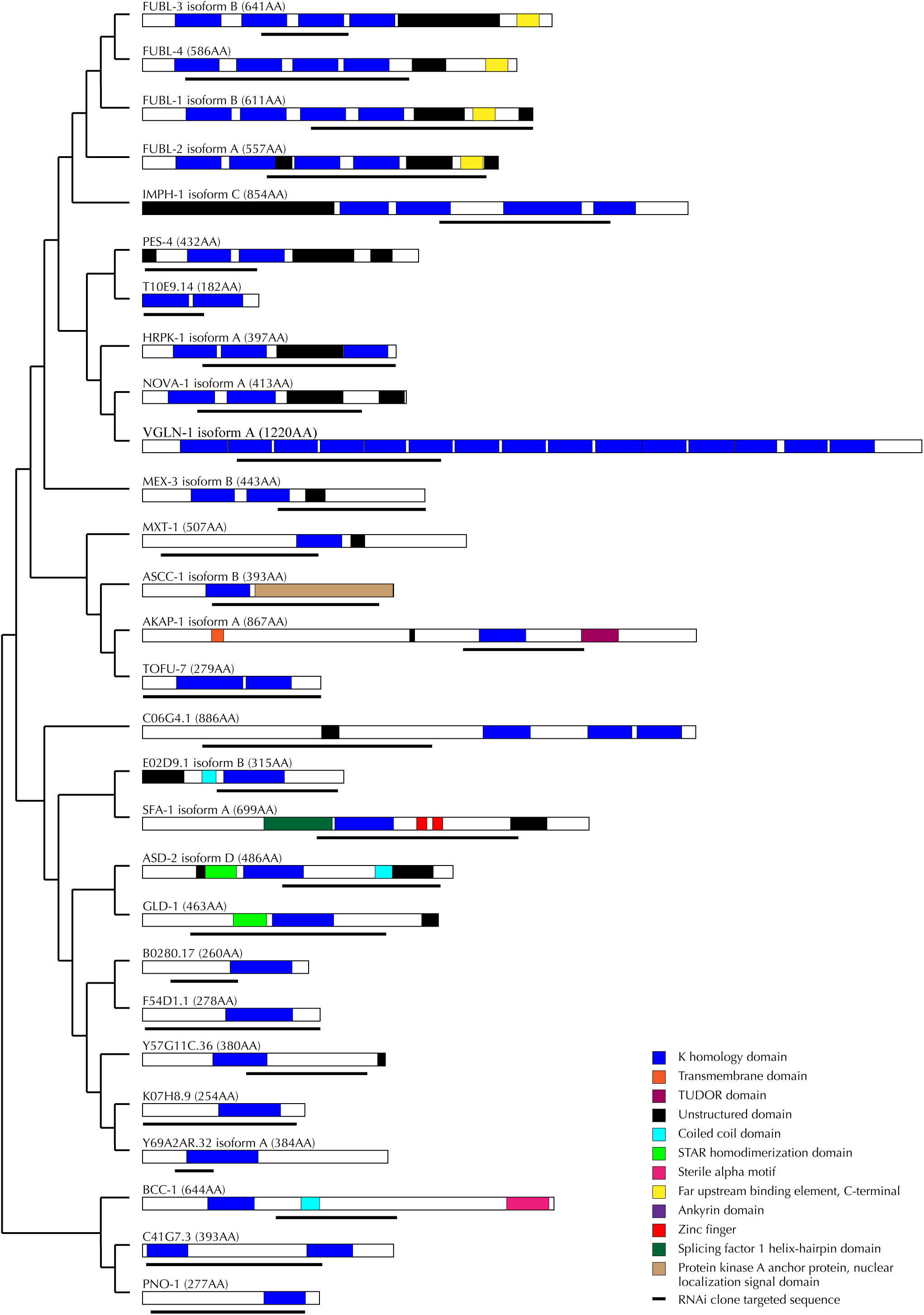
Domain architecture analysis of KH domain containing RNA-binding proteins. Protein domains prediction analysis was performed on the longest predicted isoform using SMART (Letunic and Bork 2017). Region of each protein targeted by RNAi are highlighted below the predicted protein structure. Proteins are grouped in the clades identified via our phylogenetic analysis.

## MATERIALS AND METHODS

### Worm strains

Worm culture and maintenance was performed as previously described (Brenner, 1974). Bristol N2 was used as the wildtype strain. Strains used in this study are OH3646 *lsy-6(ot150)*; *otIs114 [Plim-6-gfp + rol-6(su1006)]*, OH812 *otIs114 [Plim-p-gfp + rol-6(su1006)]*, PS3662 *syIs63 [cog-1::gfp + unc-119(+)]*, OH7310 *otIs193 [Pcog-1::lsy-6 + rol-6(su1006)]; sy1s63*, VT1296 *mir-48 mir-241(nDf51) col-19::gfp (maIs105)*, MT7626 *let-7(n2853*), VT2223 (*lin-31(n1053*); *col-19::gfp(maIs105*); *alg-1(ma202*). All strains were grown at 20° C with the exception of MT7626 *let-7(n2853*) which was grown and maintained at 15° C to prevent excess bursting.

### RNA interference

RNAi constructs (pL4440) were obtained from the Ahringer RNAi library (Kamath *et al.* 2000); Source Biosciences) except for *bcc-1* and *E02D9.1* which were obtained from the Vidal RNAi library (Rual *et al.* 2004); Source Biosciences). In addition, 3 RNAi clones were constructed by genomic amplification of the endogenous loci and cloning of the fragment into the L4440 vector. The *fubl-3* clone was generated by using forward 5’-GCCCACTAGTGGACTAACTGCAACGTTCAA-3’ and reverse 5’-GTGGGTACCATTTGCCGCCTCAGAATTG-3’. The *Y6A2AR.32* clone was generated using forward 5’-GCTCAGATCTTGCCACGTTTCATGCGAAAC-3’ and reverse 5’-GTAGGTACCGGAAGCTCTTCCTCTCACAA-3’. The *B0280.17* clone was generated using forward 5’-GGCCAGATCTCTTCTAGTTCGTGAAATCAA-3’ and reverse 5’-ATAGGTACCGCAGTCTCGGGAGGAAAG-3’. The amplified genomic fragments containing restriction sites were digested using SpeI and Kpn1 (*fubl-3*) or BglII and KpnI (*Y69A2AR.32 and B0280.17*) and were ligated with the digested L4440 vector using NEB (M2200) Quick Ligation protocol. Ligated plasmid was then transformed into *E. coli* HT115 bacteria. Sequence insertion into the L4440 plasmid was confirmed via Sanger sequencing (using M13 forward sequencing primer). Although the Ahringer clone targeting *mex-3* was obtained, RNAi of *mex-3* in *lsy-6(ot150)* and *mir-48 mir-241(nDf51)* resulted in highly penetrant embryonic lethality preventing scoring of the F1 progeny of the RNAi treated animals.

RNAi experiments were done by feeding and performed at 20° C unless otherwise stated and as described below. RNAi plates were prepared and seeded using standard methods (Kamath *et al.* 2000). Scoring requiring fluorescence was done on a Leica DM6B fluorescent compound microscope. Imaging of fluorescence-based phenotypes was done using the Leica DM6B mounted camera and processed using Leica software. Photoplates were assembled using Adobe Illustrator. Scoring of vulval bursting, brood size, and embryonic lethality were done a standard Leica dissecting microscope.

### ASEL cell fate differentiation

*Plim-6::gfp (otIs114)* and *lsy-6(ot150)*; *Plim-6::gfp (otIs114)* worms were placed on RNAi as embryos and F1 progeny were scored as L4 or young adults to increase the ease of detecting fluorescent signal in ASEL neurons. Each group of genes was scored alongside the negative control (empty L4440 vector) and our positive control (*dcr-1* RNAi). Worms were scored as cell fate defective when *lim-6::gfp* was absent in the ASEL neuron soma.

### Uterine *cog-1::gfp*

*cog-1::gfp (syIs63)* and *cog-1::gfp (syIs63); otIs193[Pcog-1::lsy-6*; *rol-6(su1006)]* worms were placed on RNAi as embryos and F1 progeny were scored at mid-late L4s in order to ensure a strong GFP signal in both vulval and uterine cells. Each group of genes was scored alongside the negative control (empty L4440 vector) and our positive control (*dcr-1* RNAi). Worms were considered to have abnormal uterine *cog-1::gfp* if either the anterior or posterior or both uterine cells were lacking GFP. *cog-1* expression was scored as normal when GFP expression was observed in both uterine cells and in vulval cells.

### Hypodermal *col-19::gfp* expression and seam cell number

*col-19::gfp (maIs105)* and *mir-48 mir-241(nDf51) col-19::gfp (maIs105)* animals were placed on RNAi as young L4s and their F1 progeny were scored as young adults. Each group of genes was scored alongside the negative control (empty L4440 vector) and our positive control (*dcr-1* RNAi). Worms were scored first for the presence of *col-19::gfp* in the hypodermal cells. Normal expression was defined as all hypodermal cells expressing *col-19::gfp* while abnormal expression was defined as GFP signal absent in many or all of hypodermal cells. Worms were also scored for the number of seam cells present between the pharynx and rectal cells; seam cells were identified using the *col-19::gfp* transgene. *lin-31(n1053)*; *col-19::gfp* (*maIs105)*; *alg-1(ma202)* worms were scored in an identical manner when assaying hypodermal *col-19::gfp* expression.

### Vulval bursting

*let-7(n2853*) worms were grown and maintained at 15°C. Embryos were synchronized by hypochloride/ NaOH solution and embryos plated directly onto RNAi plates as previously described (Parry *et al.* 2007). The embryos were hatched and grown at 15°C until young adults. Worms were scored for vulval bursting ∼ 6 hours after the L4 molt to ensure all animals had reached adulthood. Each group of genes was scored alongside the negative control (empty L4440 vector) and our positive control (*dcr-1* RNAi).

### Brood size and embryonic lethality

Wildtype (N2) worms were placed on RNAi as L4s and allowed to lay embryos. When the F1 progeny reached the L4 stage, individual hermaphrodites were moved to their own RNAi plates and allowed to lay embryos for 24 hours. After 24 hours, each animal was moved to a fresh RNAi plate each day for 3 additional days. Live larvae were counted on each plate (by picking) 24 and 48 hours after the parent has been moved to ensure all larvae were counted. Dead embryos on each plate were counted 48 hours after removal of the parent. The total number of live larvae and dead embryos for each hermaphrodite was tallied and together encompass brood size. Embryonic lethality was calculated as (# dead embryos/total brood size) x 100%. Larval arrest was rarely seen, but when it did occur these worms were counted as “live larvae” because they had successfully hatched and developed beyond the embryonic stage.

### Phylogenetic Analysis

Full proteins sequences of the longest isoforms for each protein were entered collected from Wormbase and entered to the Mega X program. A MUSCLE protein alignment was carried out to provide input for further phylogenetic analysis. In order to construct the tree, we selected the Maximum Likelihood method and bootstrapped the tree-building (1000 iterations) to increase the stringency of the method. A simple LG model was selected for the substitution model, utilizing a Nearest-Neighbor-Interchange (NNI) method. The phylogenic tree shown represents 27 of the 28 KH domain proteins in the *C. elegans* genome: *mask-1* was excluded due to extensive length and sequence/domain variability from the rest of the protein family.

### Protein domain and architecture

To generate the protein domain graphics, we first determined the longest isoform of each individual protein. The amino acid sequence of the proteins were obtained from Wormbase.org and entered into Simple Modular Architecture Research Tool (SMART) (Letunic and Bork 2017) under the Genomic options. Domain start and end points were noted and used to generate the proteins graphics in Adobe Illustrator.

To generate the coverage of each RNAi clone used in this study, primer pairs were obtained from the Ahringer library database, aligned to the appropriate transcript. Each RNAi target was then translated in the appropriate frame and aligned to the complete protein sequence. Predicted NLS sites were determined using cNLS Mapper using a threshold of 5.0 (Kosugi *et al.* 2009). Only high confidence (score > 8.0) NLS regions were included in the domain graphics.

### Data Availability

Strains and plasmids are available upon request. All data necessary for confirming the findings of this article are present within the article and the associated figures, and tables.

## DISCUSSION

### KH-domain containing RBPs functionally interact with multiple miRNA families

To determine whether *C. elegans* KH domain-containing RBPs may function with miRNAs to regulate gene expression, we asked whether RNAi knockdown of KH domain genes could modify the phenotypes observed in reduction-of-function miRNA or family mutants. Surprisingly, 19 of the 24 tested genes genetically interacted with at least one miRNA mutant background, suggesting widespread functional interaction between KH RBPs and miRNAs. Interestingly, the KH domain genes fell into two groups: those that modified phenotypes of all miRNA sensitized backgrounds tested and those that genetically interacted with specific miRNA reduction-of-function mutants (Table 1). *sfa-1 and asd-2* functionally interacted with multiple miRNA families (Table 1), suggesting that these two genes have broad roles in regulation of gene expression. The human ortholog of *sfa-1*, SF1 (Splicing Factor 1), participates in the spliceosome assembly by binding 3’ branch sites of pre-mRNAs while its partner, U2AF, cooperatively binds the 5’ branch site (Rino *et al.* 2008). Likewise, the ortholog of *asd-2, quaking*, has established roles in RNA processing, including alternative splicing and generation of select miRNAs and circular RNAs (Darbelli and Richard 2016). In *C. elegans*, both *sfa-1* and *asd-2* are predicted to play a role in splicing, with *asd-2* modulating the alternative splicing of *unc-60* and other transcripts (Kuroyanagi 2013) and *sfa-1* regulating the pre-mRNA splicing of multiple genes (Heintz *et al.* 2017). Depletion of either *sfa-1* or *asd-2* was sufficient to induce embryonic lethality and reduce brood sizes (Table 2) (Ma and Horvitz 2009; Chu *et al.* 2014), consistent with their essential roles as potential global regulators of splicing. Unbiased reverse genetic screens have previously identified splicing machinery members as important for miRNA-mediated gene regulations (Parry *et al.* 2007). Similarly, factors involved in mRNA processing, including splicing, were found to modulate RNAi efficacy (Kim *et al.* 2005). While splicing and small RNA (including miRNA) pathways intersect, the exact mechanisms by which this occurs remain largely unknown. Given *sfa-1* and *asd-2* potential roles in splicing, it is perhaps not surprising that these factors show broad functional interaction with miRNAs across all of our assays.

In contrast to the splicing-related factors, the majority of the KH domain-containing RBPs genetically interacted with specific miRNAs (Table 1). RNAi knockdown of *pes-4, T10E9.14*, and *mask-1* enhanced phenotypes of both *lsy-6(ot150)* (Figure 1 and Table 1) and *mir-48 mir-241(nDf51)* (Figure 3 and Table 1) mutants, suggesting a somewhat general role in gene regulation that spans multiple tissues. By contrast, *akap-1, C41G7.3*, and *pno-1*, genetically interacted with *mir-48 mir-241(nDf51)* (Figure 3 and Table 1), but not *let-7(n2853)* (Figure 4 and Table 1), suggesting a more narrow role for these KH domain genes in gene target regulation. Such functional separation can be achieved through differences in temporal expression or perhaps through distinct specificities of RBPs to target RNAs. In comparison, RNAi of *fubl-1, fubl-3, mxt-1*, and *B0280.17* genetically interacted with both *mir-48 mir-241(nDf51)* and *let-7(n2853)*, but not *lsy-6(ot150)* (Table 1). The *let-7-family* shared interactions suggest that these RBPs may have more general roles in developmental timing or may regulate broader sets of target genes. Interestingly, *fubl-1 (C12D8.1)* has been previously identified as a functional interactor of RNAi (Kim *et al.* 2005), suggesting that this gene’s activity may impact gene regulation carried out by multiple small RNA pathways.

### KH domain protein relatedness

Protein domains are conserved, structured portions of a protein that can fold and function independently. As distinct functional units of a protein, they can dictate, or add to, the overall cellular and molecular role of the protein. Evolution of protein structure and function is in part driven by addition or removal of domains through genetic recombination of domain-encoding gene sequences. To better understand the evolutionary and functional relatedness of the KH domain-containing proteins in *C. elegans* we performed a phylogenetic analysis (Figure 7). Our analysis highlights the overall diversity of these proteins, revealing low levels of similarity between many of the clades, consistent with the observation that in many cases, the proteins sequence similarity is limited to the KH domain(s). However, in contrast to the overall diversity of the proteins, we do see high degrees of relatedness in several of the clades, most notably those containing the FUBL proteins and the grouping consisting of GLD-1 and ASD-2 (Figure 7). This is not surprising given the similarity in domains and overall protein architecture (Figure 8). The phylogeny highlights several clades that functionally interact with miRNAs (Figure 7), perhaps reflecting the functional relatedness relevant to miRNA-mediated regulation of gene expression.

### Potential models for KH domain RBP and miRNA coordination

How might KH domain RBPs functionally interact with miRNA pathways to regulate gene expression? Given the evolutionary and domain architecture diversity, the KH RBPs may coordinate with miRNAs, directly or indirectly, via distinct mechanisms. These RBPs may directly affect aspects of miRNA biogenesis and function or they may indirectly intersect with miRNA pathways by affecting target mRNA processing, transport, stability, and degradation.

Proteins involved in splicing, such as SFA-1 and ASD-2, may be involved in splicing events that lead to the production of miRNA transcripts either from their independent gene loci or as part of host mRNA processing (Figure 9A). In this scenario, loss of a splicing factor’s function may reduce the amount of primary miRNA transcript produced, enhancing the reduction of function phenotypes observed in our sensitized backgrounds (Figure 9A). In addition, splicing factors may indirectly intersect with miRNA pathways by either increasing or decreasing the availability of a gene target (Figure 9B). Alternative splicing of 3’ UTRs that eliminate miRNA target sites has been recently observed (Han *et al.* 2018). Under this model, KH domain gene depletion could result in alternatively spliced mRNA isoforms that are no longer able to escape miRNA-mediated regulation (Figure 9B), enhancing the phenotypes observed in our reduction of function miRNA mutants.

**Figure 9.**
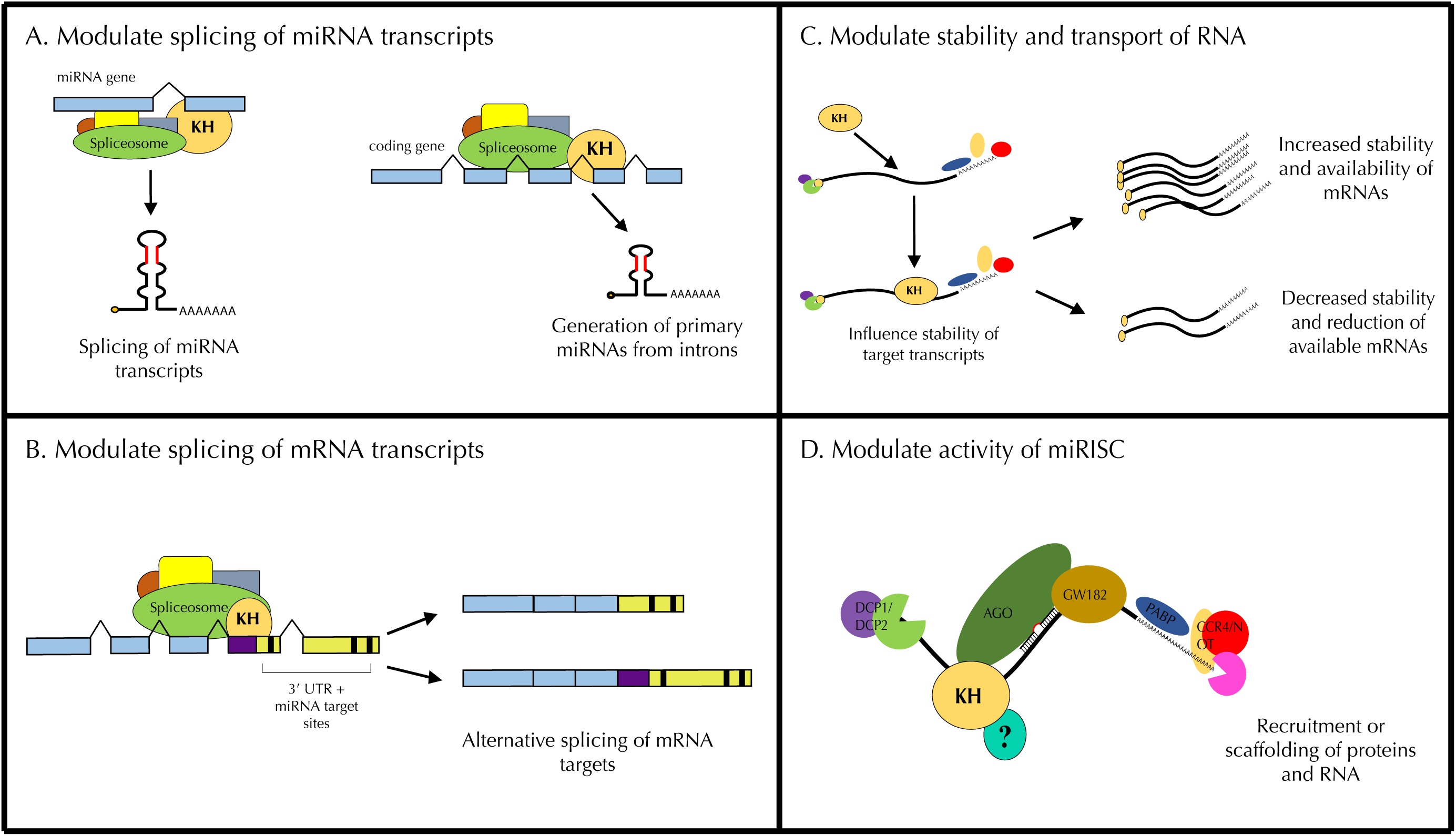
Models for potential interactions between miRNAs and KH domain-containing RNA binding proteins. (**A**) KH RBPs may modulate the splicing of primary miRNA transcripts. (**B**) RBPs my modulate the splicing of miRNA target transcripts and alter the availability of miRNA target sites in their 3’ UTRs. (**C**) RBPs may modulate the stability of miRNA target transcripts. Loss of KH domain proteins could increase the pool of target mRNAs, enhancing the miRNA reduction-of-function phenotypes, or decrease the pool of target mRNAs, resulting in the suppression of miRNA mutant phenotypes. (**D**) RBPs may modulate the activity of miRISC by bridging known RNA and protein components or by the recruitment of additional regulatory factors.

In another possible scenario, KH domain-containing factors may affect mRNA stability, localization, or transport and thus alter the pool of available miRNA targets (Figure 9C). Increased stability of target mRNAs perhaps through sequestration could reduce miRNA efficacy (Figure 9C). In contrast, reduced stability of miRNA target mRNAs could result in suppression of miRNA-related phenotypes observed in our assays. Interestingly, *Drosophila* orthologs of MXT*-*1 (MEXTLI) and B0280.17 (HOW) can enhance the stability of mRNAs (Nabel-Rosen *et al.* 2002; Hernández *et al.* 2013). The B0280.17 ortholog (HOW) shows isoform dependent enhancement or suppression of mRNA stability in order to modulate mRNA translation (Hernández *et al.* 2013). Likewise, the human orthologs of the FUBL proteins can positively or negatively modulate (depending on the protein) translation of their mRNA targets by binding the 3’ UTRs and influencing their stability (Zhang and Chen 2013). These observations lend further support to this model and suggest that the functional interactions between these RBPs could be complex and context dependent.

Lastly, it is possible that some KH domain-containing RBPs may directly interact with protein components of the miRNA pathway to modulate target gene expression. Several proteins contain additional domains that are predicted to have RNA-binding activity (SAM, zinc finger, splicing factor helix hairpin) (Figure 8) and could mediate interactions among proteins and RNA. Other functional domains such as prion-like or low complexity domains were present in approximately 50% of the RBPs tested. These domains have been implicated in driving liquid phase separation and formation of protein aggregates and RNPs (Putnam *et al.* 2019). We also see several examples of domains critical for protein-protein interactions, notably the TUDOR domain present in AKAP-1, the STAR homodimerization domains present ASD-2 and GLD-1, and the ankyrin repeats in VGLN-1. Some KH domain-containing RBPs may alter the activity of miRISC by bridging essential protein components or by recruiting additional regulatory factors (Figure 9D). This model is supported by the observation that 8 of the 29 KH domain-containing RBPs were previously found to physically interact with miRISC components or Dicer, DCR-1 (Table 3). MASK-1, FUBL-1, -2, and -3 co-precipitated with AIN-1 (Wu *et al.* 2017), while HRPK-1 and IMPH-1 co-precipitated with DCR-1 (Duchaine *et al.* 2006) and ALG-1 (Zinovyeva *et al.* 2015). GLD-1 was found to co-precipitate with ALG-1 (Akay *et al.* 2013; Zinovyeva *et al.* 2015) and AIN-2 (Zhang *et al.* 2007). These proteins may act as scaffolds for the formation of RNP complexes, bridging RNA components (mRNA or miRNA) miRNA biogenesis factors or miRISC (Figure 9D).

**Table 3.**
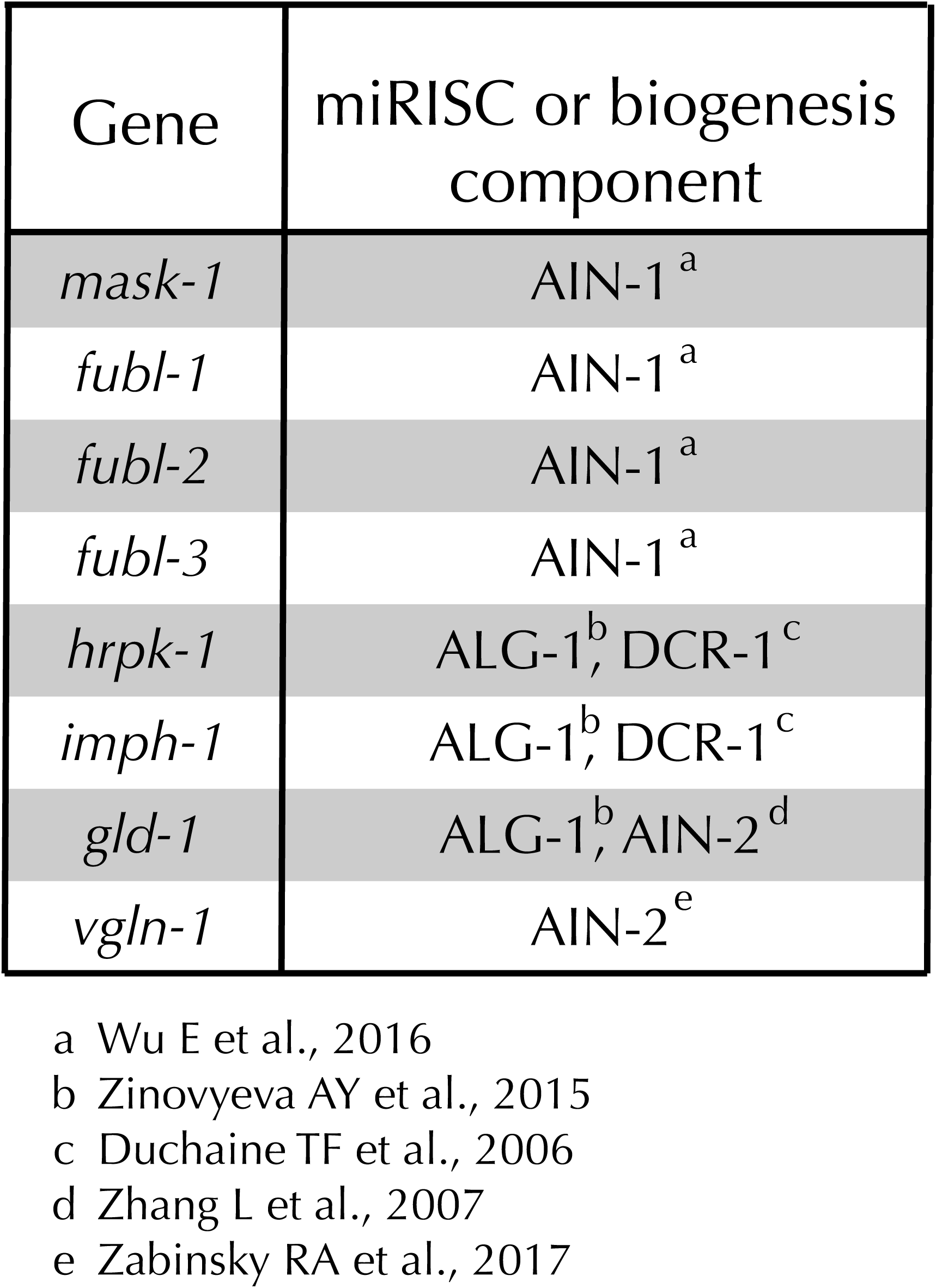
Several KH domain containing RBP’s physically interact with miRISC components

Overall, our screen demonstrated that many of the KH domain-containing RBPs in *C. elegans* functionally interact with miRNA-mediated regulation of gene expression. Further work is essential to characterize the mechanisms through which individual KH domain proteins may affect gene expression and how they might functionally intersect with miRNA pathways. This study highlights a number of candidates for future genetic, molecular, and biochemical characterization and demonstrates the extent to which miRNAs and KH domain RBPs may directly or indirectly coordinate to ultimately regulate gene expression.

## Acknowledgements

We thank members of the Zinovyeva lab for helpful technical discussions and assistance. We are grateful to Xantha Karp for critical reading of this manuscript and thank Erik Lundquist for sharing reagents. Some of the strains used in this study were provided by the CGC, which is funded by NIH Office of Research Infrastructure Programs (P40 OD010440). We thank Wormbase for providing the various necessary resources.

